# Rapid Isothermal Detection of Heavy Metals via Transcription-Factor-Gated DNA Strand Synthesis

**DOI:** 10.64898/2026.05.29.728911

**Authors:** Padric M. Garden, Alexander A. Green

## Abstract

Conventional methods for monitoring toxic heavy metals typically require sophisticated laboratory instrumentation and leave a critical gap for rapid, on-site detection. Herein we present <u>T</u>ranscription-factor-<u>O</u>ccluded <u>N</u>ick <u>E</u>xtension or TONE, a rapid, isothermal biosensing platform for heavy metal detection that exploits allosteric transcription factor (aTF) regulation to gate DNA strand-displacement amplification. In this approach, operator sequences modified with deoxyinosine (dI) substitutions are employed. When bound by an aTF, the dI sites are shielded from cleavage by Endonuclease V. Upon exposure to target heavy metals, the aTF dissociates, permitting the enzyme to nick the DNA and trigger a strand-displacement reaction. The amplified DNA is then detected via an instrument-free lateral flow assay, delivering a visual readout within 25 minutes at room temperature. We demonstrate the utility of TONE using the TetR aTF and further adapt the assay for copper and lead detection through the transcription factors CsoR and CadC, respectively, achieving detection limits as low as 40 nM and 80 nM, respectively. TONE offers a sensitive, low-cost, and field-deployable solution for environmental monitoring and other applications requiring rapid heavy metal analysis.

## INTRODUCTION

The development of biosensing approaches for the sensitive, specific, and rapid detection of small molecules and ions is critical for both fundamental research and applications such as metabolic engineering, medical diagnostics, food safety, and environmental monitoring.^1,2^ Although traditional recognition elements like enzymes, antibodies, and aptamers are well established, they often detect only a limited range of small molecules, spurring the search for new recognition elements.^1^

Allosteric transcription factors (aTFs) have emerged as a promising alternative for in vitro biosensing.^1^ Bacteria have evolved a diverse array of aTFs to sense various small molecules and ions.^3^ Typically these proteins undergo a conformational change upon ligand binding that alters their affinity for specific operator DNA sites.^4^ Although this mechanism has been widely exploited in vivo to create whole-cell biosensors (WCBs) and design genetic circuits in synthetic biology, WCBs often suffer from poor robustness, slow response times, instability, reproducibility issues, biosafety concerns, and challenges related to analyte permeability and toxicity.^1^

To overcome these limitations, recent efforts have focused on harnessing aTFs in vitro. Isolating aTFs for use in cell-free systems bypasses the complexity of living cells, enabling the development of universal, stable transduction systems that convert aTF allosteric effects into measurable signals. Building on the natural regulatory role of aTFs, platforms based on aTF-dependent in vitro transcription (IVT), such as ROSALIND^5,6^, or in vitro transcription-translation systems^7^ have proven effective. In these systems, aTFs detect small molecules and ions to regulate the transcription of RNA outputs, such as fluorescent aptamers or mRNAs encoding reporter proteins that generate signals over time. Beyond these methods, other strategies have been developed, including coupling aTF recognition with proximity driven outputs (e.g. FRET,^8,9^ fluorescence localization^10^, or electrochemical interactions^11^) and generating DNA signal outputs via enzyme competition with aTFs. Examples of these approaches include competing with DNA polymerase for G-quadruplex formation, restriction enzymes for PCR-based assays,^12^ ligase for qPCR, rolling circle amplification (RCA) and recombinase polymerase amplification (RPA),^13^ or Cas12a for collateral cleavage-based detection.^14,15^

In this work, we describe TONE, <u>T</u>ranscription-factor-<u>O</u>ccluded <u>N</u>ick <u>E</u>xtension, a novel methodology based on the competition between aTFs and Endonuclease V, an enzyme that cleaves two nucleotides 3’ of the nonstandard nucleotide deoxyinosine (dI). We demonstrate that aTFs tolerate dI substitutions within their operators and can efficiently block Endonuclease V activity when bound. The unique functionality of Endonuclease V, cleaving 3′ of the dI, provides an opportunity for aTF-dependent strand-displacement amplification (SDA), which we harness for lateral flow-based readout. Moreover, all assay components operate at room temperature, and the assay is completed in under 30 minutes, enabling a fully instrumentation-free test. We further demonstrate the ability to orthogonally detect lead and arsenic in water samples using duplex lateral flow strips.

## RESULTS

### TetR-Mediated Protection of dI-Substituted Operators

A key premise of our approach is that an allosteric transcription factor bound to its operator can occlude endonuclease-mediated cleavage (Figure 1a). We first investigated whether an aTF can tolerate deoxyinosine (dI) substitutions within its operator. To this end, we chose the well-characterized tetracycline-class antibiotic sensor TetR. While dI can pair with all four standard nucleobases, it shows a preference for pairing with dC.^16^ We likewise focused on substituting guanine residues with dI (dG→dI) to minimize disruption of the operator structure. Starting with the TetO operator sequence 5′-TCCCTATCAGTGATAGAGA-3′, we generated six possible dI-containing variants of the operator and assessed TetR binding using gel-shift assays. Under the tested conditions, we observed that introducing single dG→dI substitutions had little effect on TetR binding (Figure 1b).

**Figure 1.**
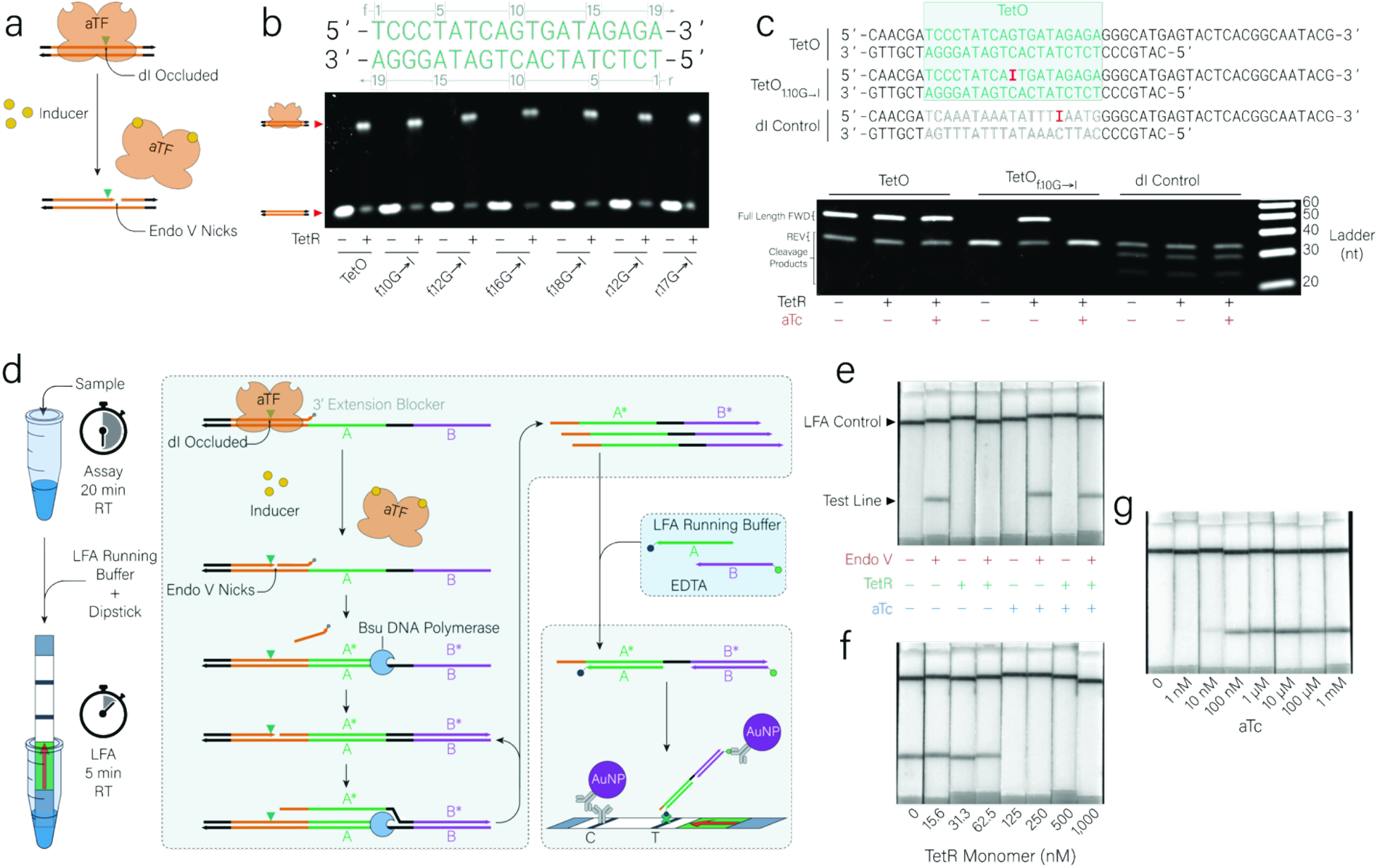
TetR-mediated protection of dI and the TONE instrument-free lateral flow assay (LFA) readout method. **a**, Incorporation of dI into the operator sequence of an allosteric transcription factor (aTF) confers protection against endonuclease V (Endo V) cleavage when the aTF is bound. Upon addition of the inducer, the aTF dissociates, permitting endonuclease V to nick the DNA two nucleotides 3’ of the dI. **b**, Gel-shift assay illustrating the impact of various dG→dI substitutions within the operator in the presence and absence of TetR. **c**, Denaturing PAGE gel demonstrating that TetR protects the dI located within the TetO operator; this protection is lost upon addition of the inducer anhydrotetracycline (aTc). **d**, Overview of the TONE assay procedure and schematic of the instrument-free assay readout mechanism. In the absence of inducer, the aTF occludes the dI, preventing endonuclease V from nicking the extension-blocked primer. Upon inducer addition, the aTF dissociates, allowing endonuclease V to nick the primer and enabling extension by Bsu DNA polymerase. Subsequent nick regeneration by endonuclease V enables amplification of the product DNA, which hybridizes with probes labeled with biotin and 6-FAM for detection on a universal LFA strip. **e**, Validation of the instrument-free assay, showing that product formation is dependent on endonuclease V, is inhibited by TetR, and is restored upon addition of the inducer aTc. **f**, Optimization of TetR concentration, indicating that 125 nM TetR is required to completely block product formation. **g**, Dose-response analysis of the assay to varying aTc concentrations, demonstrating that as little as 10 nM aTc can be detected via the formation of a visible lateral flow band.

We next examined whether TetR binding could protect dI-substituted operators from endonuclease V-mediated cleavage. Using denaturing PAGE to assess endonuclease V activity, we found that templates containing dI within the operator of TetR, when preincubated with TetR, showed no observable cleavage when endonuclease V was added (Figure 1c and Supplemental Figure 1a). These results further validate that TetR can both bind to dI-substituted operators as well as confirm they can effectively shield dI sites from endonuclease V.

### Development of an Instrument-Free Lateral Flow TONE Assay

Encouraged by these findings, we sought to translate the assay to a simple, instrument-free lateral flow format. We reasoned that nicks created by endonuclease V could serve as initiation points for a strand-displacing polymerase. Moreover, since endonuclease V nicks 3’ of a dI, the dI will remain after extension, enabling the endonuclease V to regenerate the nick. Likewise, the DNA can be repeatedly nicked and extended using a polymerase, generating product in an SDA-like manner.^17^

To implement TONE, we designed a DNA construct composed of a short forward primer annealed to a longer reverse template. The forward primer includes: (i) an anchoring domain, (ii) the operator sequence with a dI substitution, and (iii) a 3′ phosphate to block polymerase extension. The reverse template contains the complementary region to the forward primer plus the reverse complement of two probe-binding sites. In the absence of an inducer, the aTF remains bound to the operator sequence, thereby preventing endonuclease V access and blocking nick formation. When an inducer is present, the aTF dissociates from the operator, allowing endonuclease V to nick the DNA. Bsu Pol then initiates strand displacement from the nick, and endonuclease V continues to regenerate new nicking sites, leading to linear amplification of the displaced strand. The final single-stranded product is mixed with lateral flow running buffer containing probes labeled with biotin and 6-FAM, enabling detection on a universal lateral flow assay (LFA) strip (Figure 1d). This strip contains a streptavidin test line and gold nanoparticles conjugated to anti-FAM antibodies, which provides a visible readout from the amplicon-probe complex containing biotin and 6-FAM. Since TetR, Bsu Pol, and endonuclease V are all active at room temperature, this assay can be completed without the need for specialized equipment, and results can be seen within 25 minutes

To confirm that TONE product formation is specifically driven by endonuclease V-initiated nicks and modulated by TetR binding, we evaluated the combinatorial presence or absence of endonuclease V, TetR, and the TetR inducer anhydrotetracycline (aTc) (Figure 1e). In the presence of endonuclease V, nicking occurred, resulting in the generation of a detectable product. While the inclusion of TetR blocked product formation, cleavage was restored upon addition of aTc.

To maximize the TONE assay’s sensitivity for detecting low concentrations of aTc, we next sought to minimize the amount of TetR required to fully block product formation. Titration of TetR revealed that 125 nM of TetR monomer (∼62.5 nM dimer) was sufficient to completely inhibit band formation on the LFA strip (Figure 1f). Finally, we tested the lower limit of detection for aTc under these optimized conditions. Visual readout on the LFA strip showed successful detection of aTc at concentrations down to 10 nM (Figure 1g).

### Implementation of a Copper-Sensing TONE Assay

Although copper is an essential trace mineral required for numerous enzymatic and metabolic functions, elevated copper concentrations can lead to toxicity, causing severe illness or even death.^18^ To mitigate this risk, the U.S. Environmental Protection Agency (EPA) established an action limit of 1.3 ppm (∼20 *μ*M) for copper in drinking water.^19^ We hypothesized that TONE could be adapted to detect copper by employing the copper-sensing transcription factor CsoR from *B. subtilis*.^20–23^

To this end, we first confirmed that, much like TetR, CsoR could block endonuclease V activity for dI located within the CsoO operator (Figure 2a, Supplemental Figure 2a). We found that 250 nM of CsoR monomer (∼62.5 nM tetramer) was required to fully block band formation on lateral flow (Figure 2b). When first examining the responsiveness of the assay, we found that we could detect down to 100 nM of Cu^+^; however, concentrations of 100 *μ*M or higher seemed to inhibit product formation (Figure 2c).

**Figure 2.**
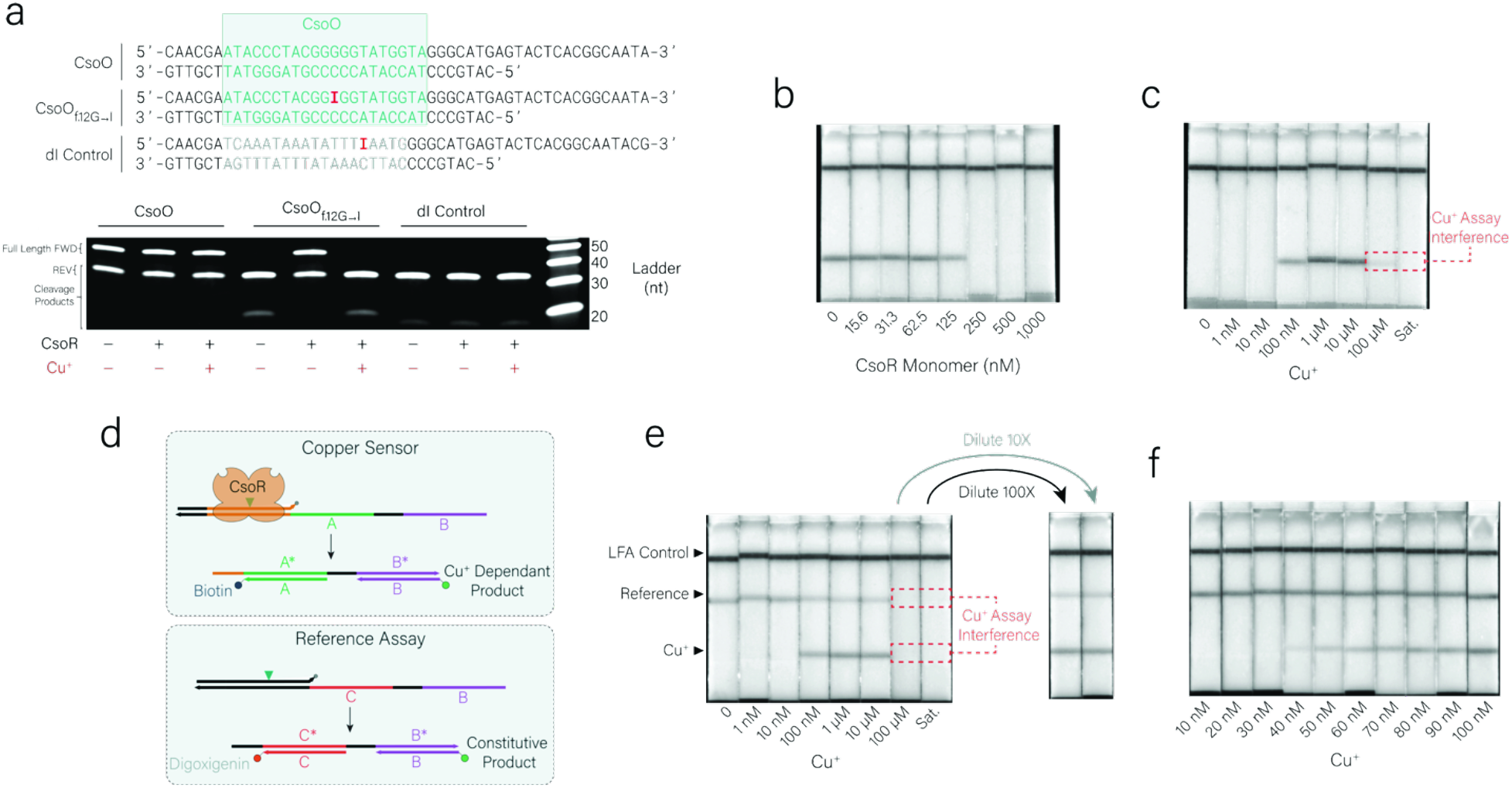
Implementation of a copper-sensing TONE assay using CsoR. **a**, Denaturing PAGE gel demonstrating that CsoR protects the dI located within the CsoO operator; this protection is lost upon addition of the inducer Cu^+^. **b**, Optimization of CsoR concentration, indicating that 250 nM CsoR is required to completely block product formation. **c**, Dose-response analysis of the assay to varying Cu^+^ concentrations, demonstrating that as little as 100 nM Cu^+^ can be detected via the formation of a lateral flow band. A loss of product is seen with 100 *μ*M and saturated Cu^+^ samples due to interference with the assay. **d**, Mechanism of the reference duplex assay. A second template containing a dI not within an operator is used to generate a constitutive product when the assay is functioning. The absence of this product indicates an invalid result. **e**, Implementation of the reference assay on duplex lateral flow. The upper test band corresponds to the reference probe and indicates invalid results at Cu^+^ concentration of ≥100 *μ*M. Diluting these samples enables readout of the presence of Cu^+^. **f**, Response of the assay to Cu^+^ concentrations between 10 and 100 nM, showing detection down to 40 nM.

To prevent false-negative results, we implemented a reference assay. In addition to the CsoO template, we added a reference template where the dI was not located within an operator sequence. This product should form constitutively and is designed to hybridize with a digoxigenin-labeled probe (Figure 2d). Products from the assay with orthogonal biotin- and digoxigenin-labeled probes can be used on duplex universal lateral flow strips with a streptavidin test line for the copper sensor and an anti-digoxigenin test line for the reference assay. The absence of this band indicates an invalid result and that the sample should be diluted before running the assay again.

Subsequent evaluation of the updated assay format demonstrated that at Cu+ concentrations of ≥100 *μ*M, both test and reference bands were absent, indicating an invalid result. However, further dilution of these samples restored assay functionality, enabling detection of higher copper concentrations (Figure 2e). We also examined the sensitivity of the system in more detail (Figure 2f) and found we could detect down to 40 nM of Cu^+^.

Cross reactivity was observed towards Cu^2+^, Zn^2+^ and Ni^2+^ (Supplemental Figure 4a). CsoR is known to interact with Zn^2+^ and Ni^2+^,^5,20^ while Cu^2+^ reactivity is likely due to some residual dithiothreitol from the enzyme storage buffers, which has previously been observed to reduce Cu^2+^ such that it can derepress CsoR.^5^

### Implementation of a Lead-Sensing TONE Assay

Lead is a naturally occurring heavy metal with no known physiological role. Low-level exposure can cause severe toxicity, including neurological impairments, developmental delays, and renal dysfunction.^24^ The U.S. EPA has established an action limit of 15 parts per billion (ppb) for lead in drinking water (approximately 72 nM). Like with copper, we hypothesized that our assay could be adapted for lead by employing the lead-and cadmium-responsive transcription factor CadC from *S. Aureus*.^25–27^

While we confirmed that CadC could protect dI within its operator (Supplemental 3b), we observed that when a dI was placed within the CadO operator, even in the absence of CadC, little product was formed in the lateral flow assay (Figure 3a). We hypothesized that this may be due to the AT-rich nature of CadO, which may prevent it from acting as an efficient primer once nicked. CadO, 5’-TACACTCAAATAAATATTTGAATGAA-3’, has a GC content of only 19%, and the G→I substitution likely further destabilizes the hybridization and ability to act as an effective primer for the polymerase.

**Figure 3.**
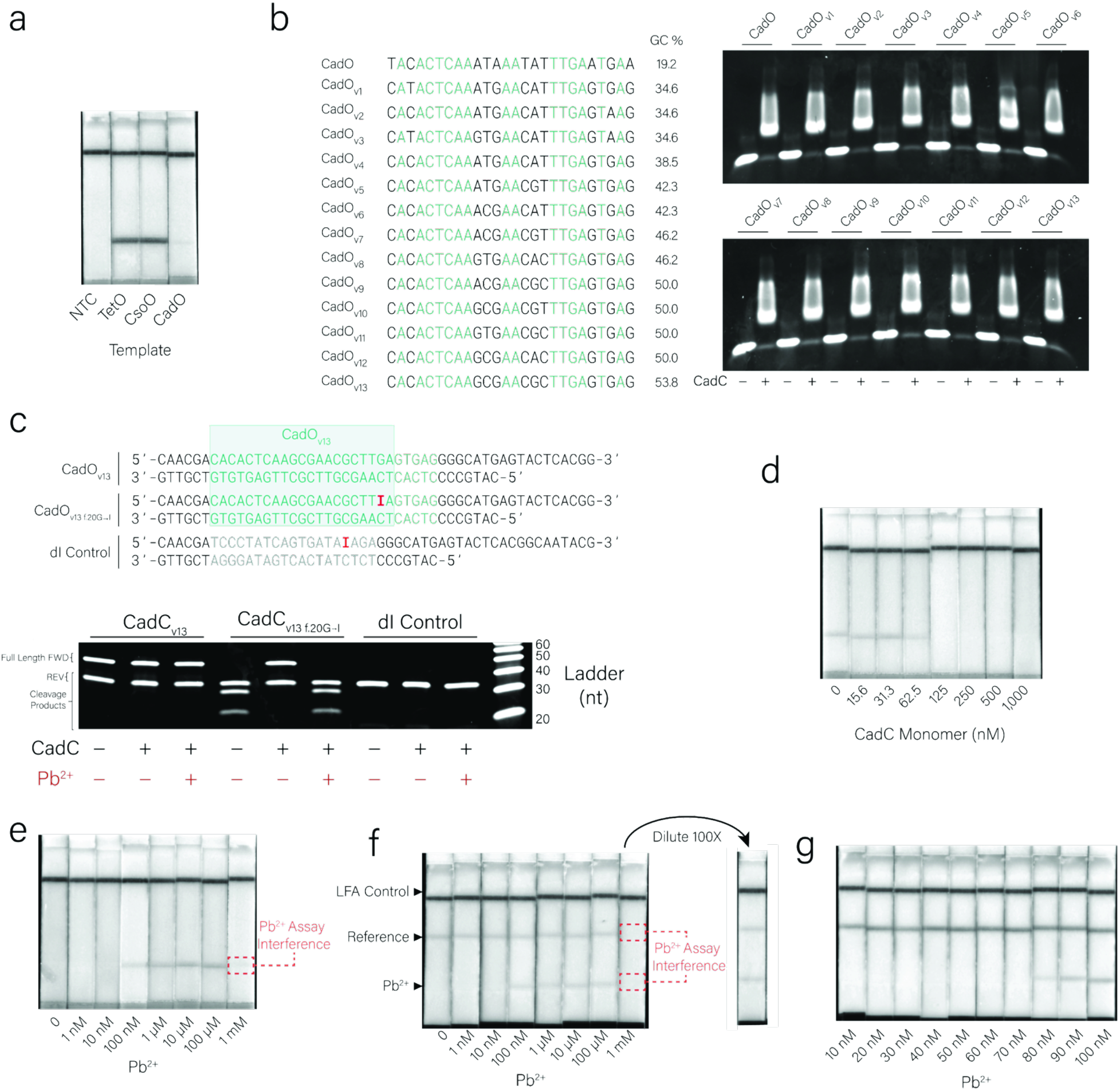
Implementation of a lead-sensing TONE assay using CadC. **a**, Lateral flow outputs of assays run without aTFs present. CadO Template produces very little product. **b**, Gel-shift assay showing the effect of increasing the GC content of the CadO operator for 13 operator variations. **c**, Denaturing PAGE demonstrating that CadC protects the dI located within the CsoO_v13_ operator; this protection is lost upon addition of the inducer Pb^2+^. **d**, Optimization of CsoR concentration, indicating that 125 nM CadC is required to completely block product formation. **e**, Dose-response analysis of the assay to varying aTc concentrations, demonstrating that as little as 100 nM Pb^2+^ can be detected via the formation of a faint lateral flow band. A loss of product is seen at 1 mM Pb^2+^ due to interference with the assay. **f**, Implementation of the reference assay on duplex lateral flow. The upper test band corresponds to the reference probe and indicates invalid results at Pb^2+^ concentration of ≥1 mM. Diluting these samples enables readout of the presence of Pb^2+^. **g**, Response of the assay to Pb^2+^ concentrations between 10 and 100 nM, showing detection down to 80 nM.

To combat this issue, we attempted to increase the GC content of the CadO operator without interfering with the ability of CadC to bind. We designed 13 variants of the CadO operator maintaining highly conserved bases while attempting to maximize the GC content of the operator. CadC tolerated these substitutions well and we successfully increased the GC content of the operator to 54% (CadO_v13_) without impacting CadC’s binding (Figure 3b). Using the CadO_v13_ operator we then demonstrated the ability of CadC to block endonuclease V activity (Figure 3c).

This new template was able to generate sufficient signal on lateral flow and we found that 125 nM of CadC monomer (∼62.5 nM dimer) was required to fully block band formation on lateral flow (Figure 3d). When examining the responsiveness of this TONE assay towards Pb^2+^, we found that we could detect down to 100 nM; however, concentrations of 1mM or higher seemed to inhibit product formation (Figure 2e).

Like with CsoR, we implemented a reference assay, demonstrating that at Pb^2+^ concentrations of ≥1 mM, both test and reference bands were absent, indicating an invalid result. However, further dilution of a 1 mM sample restored assay functionality, enabling accurate detection of higher lead concentrations (Figure 2f).

We also examined the sensitivity of TONE in more detail (Figure 2g) and found we could detect down to 80 nM of Pb^2+^. As expected, strong cross reactivity was observed towards Cd^2+^, and slight activation was also observed with Zn^2+^ (Supplemental Figure 4b). CadC is known to interact with Zn^2+^,^28–31^ and derepression with Zn^2+^ has previously been observed in vitro.^5^

### Multiplex Metal Detection on Lateral Flow

With the system able to detect two different metal ions, we next sought to implement the assays together to have a multiplex readout. First, we modified the assay such that the CsoR product binds a digoxigenin-labeled probe, the CadC product binds a biotin-labeled probe, and the reference assay binds a Cy5-labeled probe (Figure 4a). We can then detect each output on a lateral flow with anti-digoxigenin, anti-Cy5, and streptavidin test lines (Figure 4b). Using these multiplex assays, we achieved similar sensitivities for each of the assays compared to when they were run separately, and the reference assay retained the ability to indicate interference due to high metal concentrations (Figure 4c). Lastly, we confirmed orthogonal detection of both Cu^+^ and Pb^2+^ and examined the functionality of the system in complex water samples (Figure 4d). We observed no interference between the Cu^+^ and Pb^2+^ assays and the multiplex assay functioned in both tap water and river water samples, indicating that this assay could be applied to test for heavy metals in drinking water and ecological water samples.

**Figure 4.**
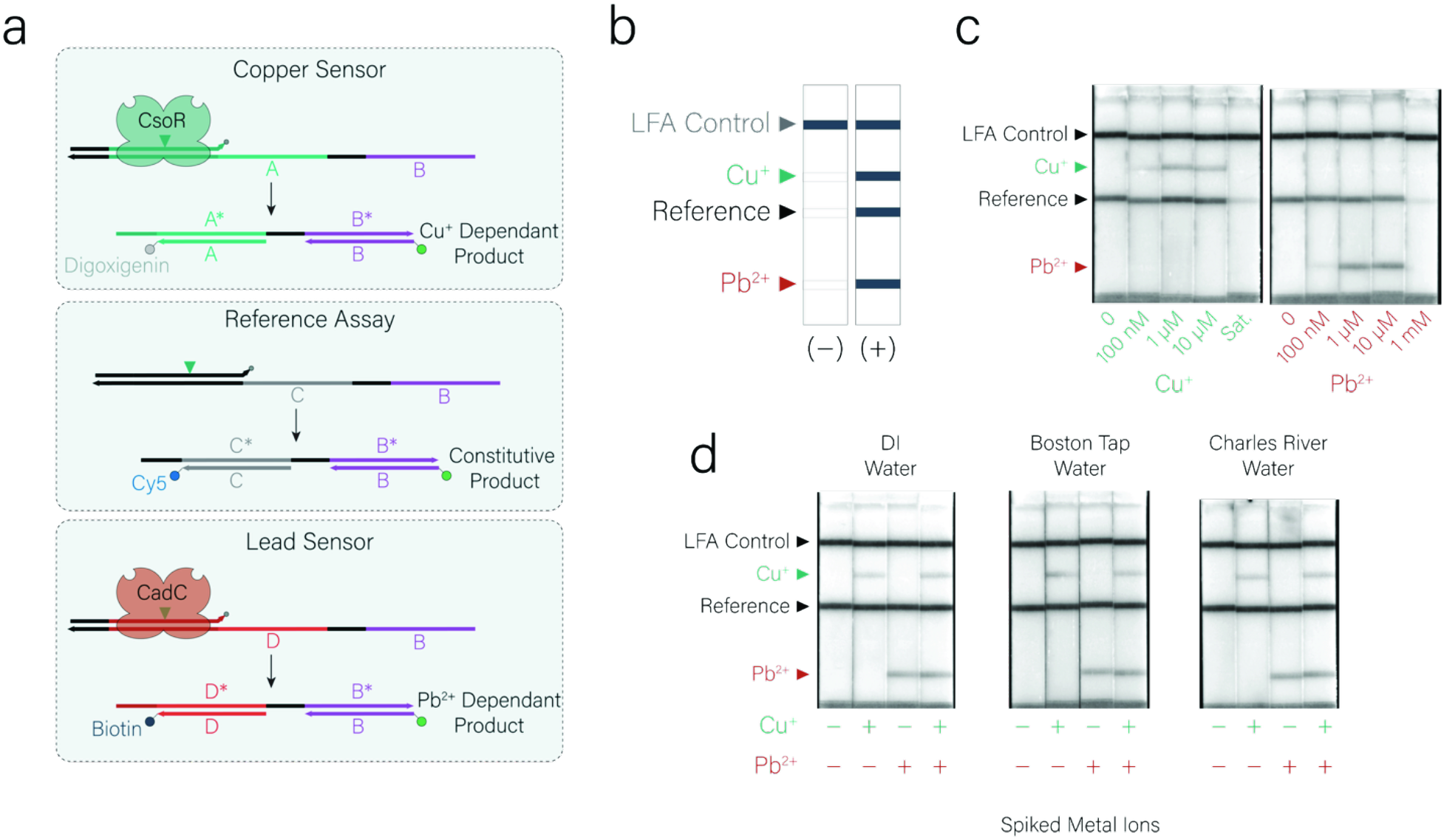
Multiplex detection of metal ions using TONE. **a**, Design of the multiplex assay. The copper-sensing assay produces a product that will hybridize to a digoxigenin-labeled probe. The reference assay binds a Cy5-labeled probe, while the lead sensor binds a biotin-labeled probe. These can be used to produce signals on a multiplex lateral flow strip with anti-digoxigenin, anti-cy5, and streptavidin test lines. **b**, Organization of the test lines on the multiplex lateral flow strip. **c**, Response of the multiplex system to Cu^+^ and Pb^2+^ confirming the functionality of each assay and the ability of the reference assay to indicate interference. **d**, Orthogonal detection of Cu^+^ and Pb^2+^ in spiked deionized water, Boston tap water, and Charles River water to demonstrate functionality in complex sample matrices.

## DISCUSSION

In this work, we introduced a rapid, isothermal, and instrumentation-free strategy for detecting heavy metals based on steric competition between allosteric transcription factors (aTFs) and endonuclease V. We showed that aTFs (TetR, CsoR, and CadC) can tolerate single dI substitutions in their operator sequences. When these aTFs are bound, they prevent endonuclease V from nicking the DNA. Upon addition of an inducer, the aTF dissociates, allowing endonuclease V to generate nicks that initiate a strand-displacement amplification at room temperature. These amplification products can then be quickly detected via a lateral flow assay, providing a convenient visual readout within 25 minutes. We demonstrated the adaptability of this method by detecting nanomolar concentrations of copper and lead ions, underscoring its potential for straightforward, point-of-use environmental testing and water-quality monitoring. Moreover, we demonstrated that this technique can be multiplexed through the use of multiple aTFs and templates to provide a rapid readout of multiple metals and can function in spiked tap water and river water samples.

Placed alongside other aTF-based detection methods, our approach is distinct in offering a fully instrument-free assay readout by employing assay components that all function at room temperature and the use of a lateral flow readout that does not require any specialized equipment. Additionally, our system relies solely on DNA rather than RNA, and it thus avoids RNase interference. Moreover, the assay appears to exhibit substantially improved sensitivity than some previous aTF-based detection systems using the same aTFs,^5^ while IVT-based platforms with engineered aTFs have demonstrated sensitivities similar to ours (50 nM Pb^2+^).^32^

Due to the mechanism we employ, our system is limited to aTFs that have binding/unbinding allosteric mechanisms. We cannot utilize classes of transcription factors that instead alter promoter architecture or interact with RNA Polymerase. For example, MerR family TFs have attractive specificity for heavy metals, but they remain bound regardless of inducer concentration and function by rearranging promoter spacing, making them incompatible with our current design.^33^

Our system is sensitive to high concentrations of metal ions, which can interfere with biochemical processes. Although the inclusion of a reference assay partially mitigates this interference, it cannot differentiate between various interfering metal ions in complex samples. Moreover, many allosteric transcription factors (aTFs) exhibit cross-reactivity with different sensitivities to various analytes, complicating the accurate determination of analyte concentrations. Combinatorial RNA logic based on the outputs of multiple aTF-dependent in vitro transcription (IVT) reactions has been demonstrated previously.^5,6^ A similar approach employing DNA logic circuits could enhance specificity by integrating responses from multiple transcription factors.

Additionally, nucleases present in complex samples may degrade assay components, an issue that could potentially be addressed through protective modifications—such as methylation, 3’ exonuclease blockers, and phosphorothioate bonds.^34^ However, protecting DNA products from nucleases remains challenging. Encapsulation of ion-sensing IVT systems in vesicles has also been demonstrated,^35^ and this strategy could be applied to our system to safeguard assay components from degradation. Finally, heavy metal efflux pumps could potentially be used^36^ to facilitate the selective transfer of ions, preventing larger molecules from entering vesicles or even concentrating low-abundance ions.

The aTFs we have tested so far are tolerant of single dG→dI substitutions, indicating this approach should be easily adaptable to additional aTFs. aTFs for new targets can be mined^11,37^ or engineered,^4,38^ to expand the possible targets of aTF-based sensors, enabling sensing of small molecules and ions that are difficult to detect by conventional biosensing approaches. Additionally, sensitivity and specificity of existing aTFs can be improved through mutagenesis or rational design.^32,39^ Future work could focus on improving specificity and thresholding via DNA logic circuits and protective vesicle-based architectures, as well as engineering or discovering new aTFs for diverse analytes. These advances would help transform this platform into an even more versatile technology for on-site monitoring of an expanding range of contaminants, paving the way for more reliable, field-deployable diagnostic and environmental surveillance tools.

## METHODS

### Materials

DNA oligonucleotides were purchased from IDT (see Supplemental Table 1 for sequences). Protein expression plasmids were purchased from Twist Biosciences. NiCo21(DE3) Competent *E. coli*, endonuclease V, Bsu DNA Polymerase, dNTPs, ET SSB, and NEBuffer r2.1 were purchased from New England Biolabs. HiTrap TALON crude columns were purchased from Cytiva and run on an AKTA Go FPLC. MagicMedia autoinduction media and 10X Native Agarose Gel loading dye were purchased from ThermoFisher Scientific. PAGE gels and Urea Loading Dye were purchased from BioRad. BugBuster 10X Protein Extraction Reagent, cOmplete EDTA-free Protease Inhibitor Cocktail, Lysozyme, Salt Active Nuclease, LB media, anhydrotetracycline, copper(I) chloride, copper (II) sulfate pentahydrate, lead (II) chloride and cadmium chloride were purchased from Millipore Sigma. TBS, imidazole, and phosphate buffers were purchased from Boston Bioproducts. Lateral flow strips were purchased from Milenia Biotec.

### aTF Protein Expression

NiCo21 cells were transformed according to the manufacturer’s protocol using 1 *μ*L of a 10 ng/*μ*L expression plasmid prior to plating. After overnight incubation, single colonies were selected and grown in 10 mL LB medium at 37°C and 300 rpm. Next, 1 mL of the overnight culture was transferred to 200 mL of Magic Media in a 1 L baffled flask and incubated at 37°C and 300 rpm for 24 hours. Cell pellets were harvested by repeated centrifugation (10 minutes at 4,000 g, 4°C) and stored at -80°C until further use.

### FPLC Buffer Preparation

FPLC extraction buffer was prepared to contain 50 mM sodium phosphate, 500 mM NaCl, and 10% glycerol (pH 7.4). FPLC wash and elution buffers were similarly prepared, except with 10 mM and 150 mM Imidazole, respectively.

### Lysate Preparation

Frozen cell pellets derived from 200 mL of Magic Media Expression were supplemented with one tablet of cOmplete EDTA-free Protease Inhibitor Cocktail. The cells were resuspended in an extraction buffer consisting of 18 mL FPLC extraction buffer, 2 mL BugBuster 10X Protein Extraction Reagent, and 200 *μ*L 1 M magnesium acetate. To the suspension, 400 *μ*L lysozyme (10 mg/mL), 2 *μ*L Benzonase, and 5 *μ*L Salt Active Nuclease were added, and the mixture was incubated on a mixer for 20 minutes. Then, 20 mL of Talon crude wash buffer was added, and the suspension was gently inverted to mix. After centrifugation at 4,100 g for 15 minutes, the supernatant was sequentially filtered through 1 *μ*m and 0.45 *μ*m syringe filters, followed by vacuum filtration through a 0.22 *μ*m filter into a 50 mL tube.

### FPLC Purification

After the FPLC system was primed and equilibrated with FPLC wash buffer, the prepared lysate was loaded onto the column, followed by a wash with 25 mL of FPLC wash buffer before elution with FPLC elution buffer. 12 fractions centered on the fraction with the highest UV absorbance were run on an SDS-PAGE and those with a purity of >95% were pooled. Pooled samples were concentrated and buffer exchanged into 1X TBS using a Pierce PES 3K MWCO filter.

### Gel-Shift Assays

50 nM duplex operator DNA was mixed with 1 *μ*M of aTF (or buffer control) in 1X TBS for 30 minutes at RT. 10X Native Agarose Loading dye was added to a final concentration of 1X before samples were loading onto 5% pre-cast BioRad TBE Native PAGE Gels, which were run at 180 V for 35 minutes before staining with 1X SYBR Gold in 1X TBE for 30 minutes.

### dI Protection Denaturing Nucleic Acid PAGE

50 nM duplex operator DNA was mixed with 1 *μ*M of aTF and 10 *μ*M Inducer (or buffer controls) in 1X NEBuffer r2.1 for 30 minutes at RT. 300 U/*μ*L of endonuclease V was added and the samples were incubated for 20 minutes at RT before the addition of TBE-Urea sample buffer added to a final concentration of 1X before samples were denatured for 10 minutes at 95°C. Samples were loaded onto 15% pre-cast BioRad TBE-Urea PAGE Gels, which were run at 180 V for 35 minutes before staining with 1X SYBR Gold in 1X TBE for 30 minutes.

### Instrument-Free TONE Assay

50 *μ*L assay reaction mixes consisted of 5 nM DNA template, 15 to 1,000 nM aTF, 50 U/*μ*L endonuclease V, 12 U/*μ*L Bsu Pol, 10 ng/*μ*L ET SSB, 200 *μ*M dNTPs, 30 *μ*L sample in 1X NEBuffer r2.1. After incubation for 20 minutes, 50 *μ*L LFA running buffer (1X TBS, 10mM EDTA, 1% Tween-20, 200 nM biotin-probe, 200 nM 6-FAM-probe) was added before the addition of a LFA strip. The LFA was allowed to resolve for 5 minutes before imaging.

### Instrument-Free TONE Assay with Reference

Assays were run as described above with the addition of 5 nM Reference DNA Template to the assay mix, and 200 nM of digoxigenin-probe to the LFA Running Buffer.

### Two-Plex Instrument-Free TONE Assay with Reference

50 *μ*L assay reaction mixes consisted of 5 nM CsoR template, 5 nM CadC Template, 5 nM Reference Template, 250 nM CsoR, 125 nM CadC, 50 U/*μ*L endonuclease V, 12 U/*μ*L Bsu Pol, 10 ng/*μ*L ET SSB, 200 *μ*M dNTPs, 30 *μ*L sample in 1X NEBuffer r2.1. After incubation for 20 minutes, 50 *μ*L LFA Running Buffer (1X TBS, 10mM EDTA, 1% Tween-20, 200 nM biotin-probe, 200 nM digoxigenin-probe, 200 nM Cy5-probe, 200 nM 6-FAM-probe) was added before the addition of a LFA strip. The LFA was allowed to resolve for 5 minutes before imaging.

## Author contributions

P.M.G. and A.A.G. conceived the idea. P.M.G. developed the methodology, designed, performed the experiments and data analysis. A.A.G. supervised the research and acquired funding. P.M.G. and A.A.G. wrote the manuscript.

## Acknowledgments

This work was supported by Defense Advanced Research Projects Agency (DARPA) funding (Contract No. N66001-23-2-4042) and Boston University startup funds to AAG. The views, opinions and/or findings expressed are those of the authors and should not be interpreted as representing the official views or policies of the Department of Defense or the U.S. Government. This material is based upon work supported by the National Science Foundation Graduate Research Fellowship under Grant No. DGE-1840990 to PMG. Any opinion, findings, and conclusions or recommendations expressed in this material are those of the authors(s) and do not necessarily reflect the views of the National Science Foundation.

