## Supporting Information for "Rapid Isothermal Detection of Heavy Metals via Transcription-Factor-Gated DNA Strand Synthesis"

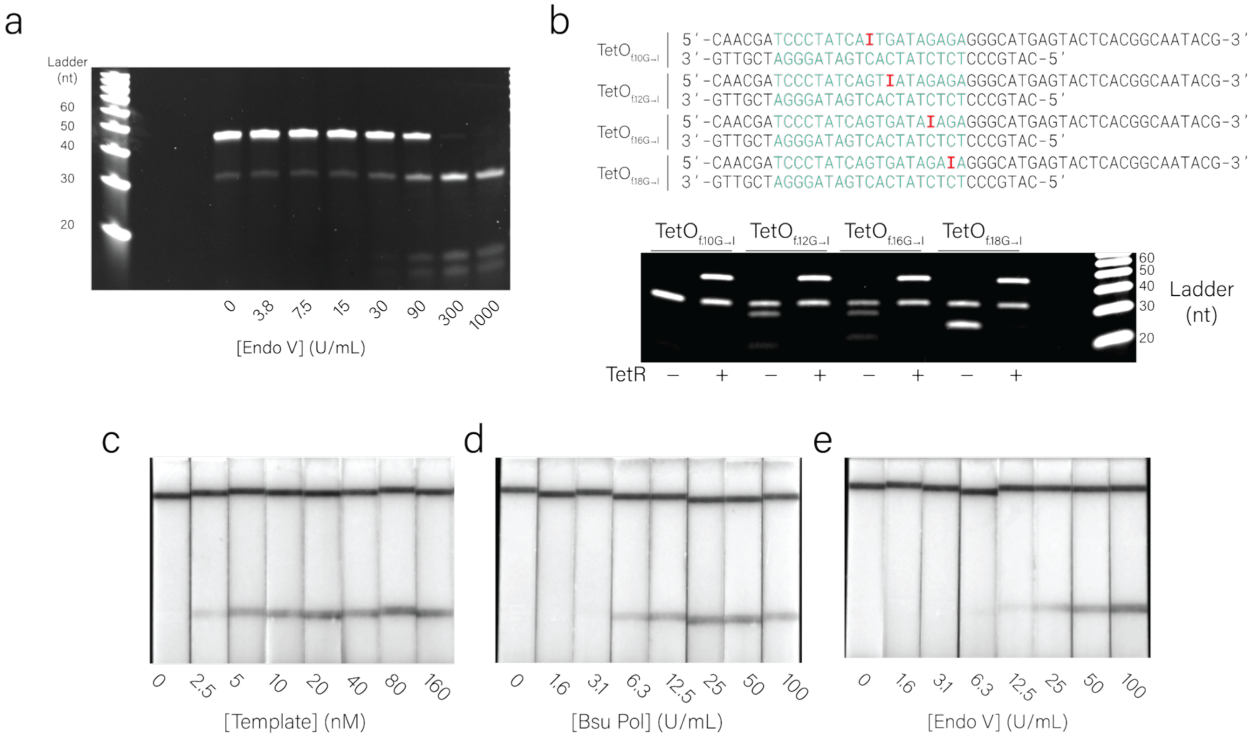


**Supplemental Figure 1. TetR dI screening and lateral flow assay optimization. a,** Optimization of endonuclease V (Endo V) concentration needed to fully cleave 50 nM of dI-containing DNA within 20 minutes at room temperature. **b,** Denaturing PAGE gel showing TetR-dependent cleavage of four possible G→I substitutions within the TetO operator. **c,** Optimization of DNA template concentration in the instrument-free LFA assay. **d,** Optimization of Bsu Pol concentration. **e,** Optimization of Endo V concentration.

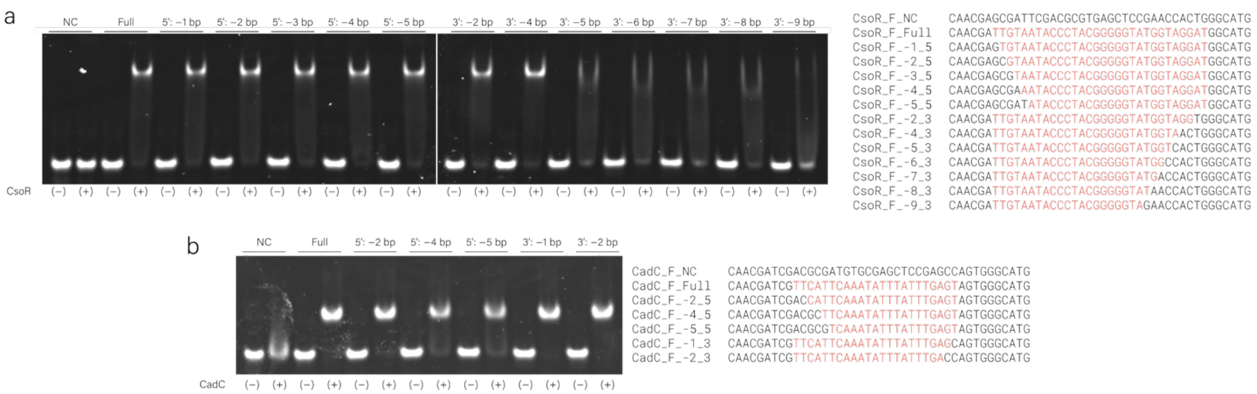

**Supplemental Figure 2. Operator Minimization for CsoR and CadC.** To determine the minimum length of the CsoR and CadC operator, gel-shift experiments were performed with truncated operators. **a,** Binding ability of CsoR to truncated CsoO operators, indicating five bases from the 5' end and four bases from the 3' end could be removed without affecting CsoR binding. **b,** Binding ability of CadC to truncated CadO operators, indicating two bases from the 5' end and two bases from the 3' end could be removed without affecting CadC binding.

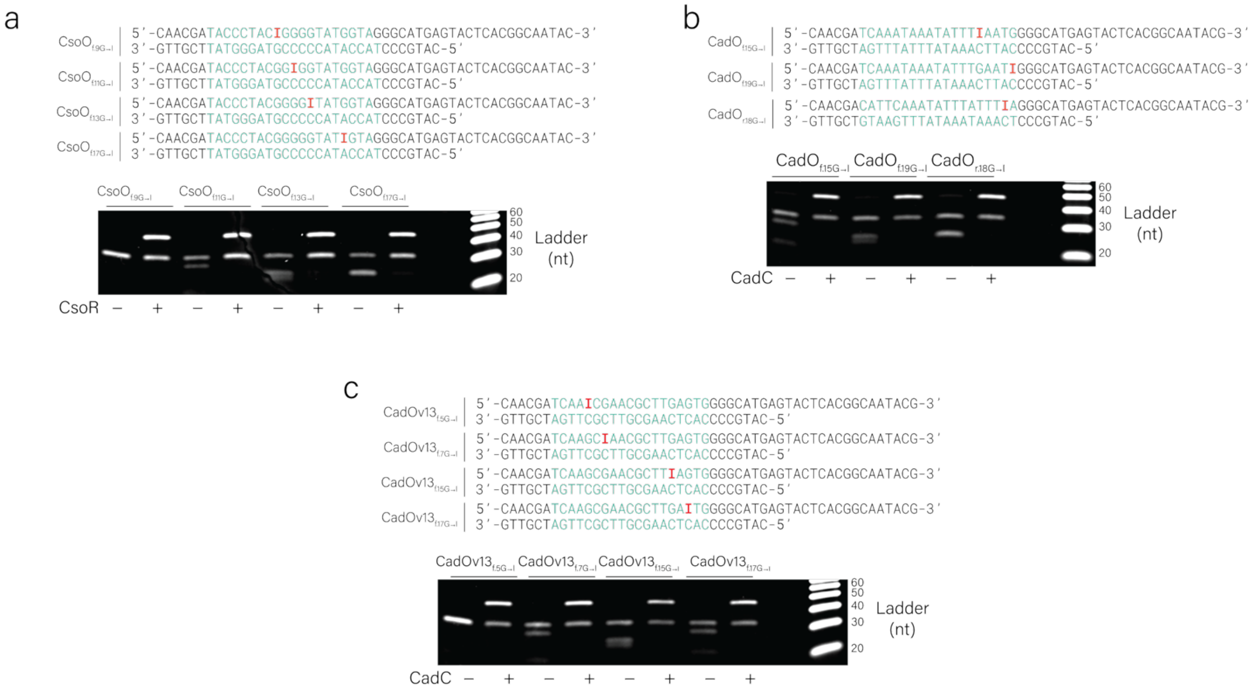

**Supplemental Figure 3. dI Screening for CsoO, CadO and CadOv13. a,** Denaturing PAGE gel showing CsoR-dependent cleavage of four possible G→I substitutions within the CsoO operator upon exposure to Endo V. **b,** Denaturing PAGE showing CadC-dependent cleavage of three possible G→I substitutions within the CadO operator upon exposure to Endo V. **c,** Denaturing

PAGE showing CadC-dependent cleavage of four possible G→I substitutions within the CadOv13 operator upon exposure to Endo V.

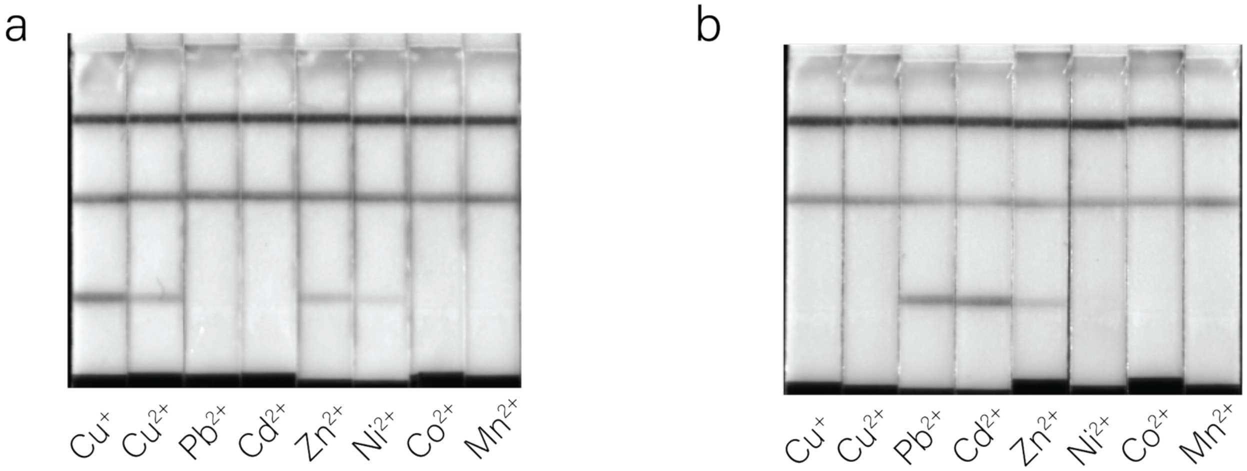


**Supplemental Figure 4. Cross reactivity of CsoR- and CadC-based assays. a,** Response of CsoR-based copper sensing assay to 10 $\mu$M of metal ions. **b,** Response of CadC-based lead sensing assay to 10 $\mu$M of metal ions.

### Supplemental Table 1: Sequences Used

| Name | Sequence |
|---|---|
| TetO_FWD_Std | CAACGATCCCTATCAGTGATAGAGAGGGCATG |
| TetO_FWD_dI1 | CAACGATCCCTATCA/ideoxyI/TGATAGAGAGGGCATG |
| TetO_FWD_dI2 | CAACGATCCCTATCAGT/ideoxyI/ATAGAGAGGGCATG |
| TetO_FWD_dI3 | CAACGATCCCTATCAGTGATA/ideoxyI/AGAGGGCATG |
| TetO_FWD_dI4 | CAACGATCCCTATCAGTGATAGA/ideoxyI/AGGGCATG |
| TetO_REV_Std | CATGCCCTCTCTATCACTGATAGGGATCGTTG |
| TetO_REV_dI1 | CATGCCCTCTCTATCACT/ideoxyI/ATAGGGATCGTTG |
| TetO_REV_dI2 | CATGCCCTCTCTATCACTGATA/ideoxyI/GGATCGTTG |
| TetO_REV_dI3 | CATGCCCTCTCTATCACTGATAG/ideoxyI/GATCGTTG |
| TetO_REV_dI4 | CATGCCCTCTCTATCACTGATAGG/ideoxyI/ATCGTTG |
| TetO_Cleavage_NC_FWD | CAACGATCCCTATCAGTGATAGAGAGGGCATGAGTACTCACGGCAATACG |
| TetO_Cleavage_REV | CATGCCCTCTCTATCACTGATAGGGATCGTTG |
| TetO_Cleavage_dI1_FWD | CAACGATCCCTATCA/ideoxyI/TGATAGAGAGGGCATGAGTACTCACGGCAATACG |
| TetO_Cleavage_dI2_FWD | CAACGATCCCTATCAGT/ideoxyI/ATAGAGAGGGCATGAGTACTCACGGCAATACG |
| TetO_Cleavage_dI3_FWD | CAACGATCCCTATCAGTGATA/ideoxyI/AGAGGGCATGAGTACTCACGGCAATACG |

| | |
|---|---|
| TetO_Cleavage_dI4_FWD | CAACGATCCCTATCAGTGATAGA/ideoxyI/AGGGCATGAGTACTCACGGCAATACG |
| dI_Control_Cleavage_FWD | CAACGATCAAATAAATATTTIAATGGGGCATGAGTACTCACGGCAATACG |
| dI_Control_Cleavage_REV | CATGCCCCATTCAAATATTTATTTGATCGTTG |
| CsoO_Cleavage_NC_FWD | CAACGATACCCTACGGGGGTATGGTAGGGCATGAGTACTCACGGCAATAC |
| CsoO_Cleavage_REV | CATGCCCTACCATACCCCCGTAGGGTATTCGTTG |
| CsoO_Cleavage_dI1_FWD | CAACGATACCCTAC/ideoxyI/GGGGTATGGTAGGGCATGAGTACTCACGGCAATAC |
| CsoO_Cleavage_dI2_FWD | CAACGATACCCTACGG/ideoxyI/GGTATGGTAGGGCATGAGTACTCACGGCAATAC |
| CsoO_Cleavage_dI3_FWD | CAACGATACCCTACGGGG/ideoxyI/TATGGTAGGGCATGAGTACTCACGGCAATAC |
| CsoO_Cleavage_dI4_FWD | CAACGATACCCTACGGGGGTAT/ideoxyI/GTAGGGCATGAGTACTCACGGCAATAC |
| CadO_Cleavage_NC_FWD | CAACGATCAAATAAATATTTGAATGGGGCATGAGTACTCACGGCAATACG |
| CadO_Cleavage_REV | CATGCCCCATTCAAATATTTATTTGATCGTTG |
| CadO_Cleavage_dI1_FWD | CAACGATCAAATAAATATTT/ideoxyI/AATGGGGCATGAGTACTCACGGCAATACG |
| CadO_Cleavage_dI2_FWD | CAACGATCAAATAAATATTTGAAT/ideoxyI/GGGCATGAGTACTCACGGCAATACG |
| CadO_Cleavage_dI3_b_FWD | CAACGACATTCAAATATTTATTT/ideoxyI/AGGGCATGAGTACTCACGGCAATACG |
| CadO_Cleavage_b_REV | CATGCCCTCAAATAAATATTTGAATGTCGTTG |
| CadOv13_Cleavage_NC_FWD | CAACGATCAAGCGAACGCTTGAGTGGGGCATGAGTACTCACGGCAATACG |
| CadOv13_Cleavage_REV | CATGCCCCACTCAAGCGTTCGCTTGATCGTTG |
| CadOv13_Cleavage_dI1_FWD | CAACGATCAA/ideoxyI/CGAACGCTTGAGTGGGGCATGAGTACTCACGGCAATACG |
| CadOv13_Cleavage_dI2_FWD | CAACGATCAAGC/ideoxyI/AACGCTTGAGTGGGGCATGAGTACTCACGGCAATACG |
| CadOv13_Cleavage_dI3_FWD | CAACGATCAAGCGAACGCTT/ideoxyI/AGTGGGGCATGAGTACTCACGGCAATACG |
| CadOv13_Cleavage_dI4_FWD | CAACGATCAAGCGAACGCTTGA/ideoxyI/TGGGGCATGAGTACTCACGGCAATACG |
| TetO_Template | GTCAGGATTAGTCCGTCAACATCGGAATATGCGAGCACCACGCCCGATAAACATGCCCTCTC TATCACTGATAGGGACGTTGCTCGCT |
| TetO_Primer | GCGAGCAACGTCCCTATCA/ideoxyI/TGATAGAGAGGGCAT/3Phos/ |
| CadOv13_Template | GTCAGGATTAGTCCGTCAACATCGGAATATGCGAGCACCACGCCCGATAAACATGCGCCAC TCAAGCGTTCGCTTGATCGTGCTCGC |
| CadOv13_Primer | GCGAGCACGATCAAGCGAACGCTT/ideoxyI/AGTGGCGCAT/3Phos/ |
| CsoO_Template_bio | GTCAGGATTAGTCCGTCAACATCGGAATATGCGAGCACCACGCCCGATAAACATGCGCTAC CATACCCCCGTAGGGTATTCGTGCTCGC |
| CsoO_Template_dig | GTCAGGATTAGTCCGTCAACATCGGAAGATATCTAATGACACATCAGGAATGATGCGCTACC ATACCCCCGTAGGGTATTCGTGCTCGC |
| CsoO_Primer | GCGAGCACGAATACCCTACGG/ideoxyI/GGTATGGTAGCGCAT/3Phos/ |
| Reference Template | GTCAGGATTAGTCCGTCAACATCGGAACGTCATTGCACAATTTGCCGATCAGTGCACTGTAG ATGCCGTTGCTCGC |
| Reference Primer | GCGAGCAACGGCATCTACA/ideoxyI/TGCA/3Phos/ |
| FAM_Capture | /56-FAM/TTTTTGTCAGGATTAGTCCGTCAACATCGG |

| | |
|---|---|
| Biotin_Capture_1 | TATGCGAGCACCACGCCCGATAAACTTTTT/3BioTEG/ |
| Biotin_Capture_2 | GATATCTAATGACACATCAGGAATGTTTTT/3BioTEG/ |
| Dig_Capture_1 | GATATCTAATGACACATCAGGAATGTTTTT/3DiG_N/ |
| Cy5_Capture_1 | CGTCATTGCACAATTTGCCGATCAGTTTTT/3Cy5Sp/ |
| CadO_WT_FWD | TACACTCAAATAAATATTTGAATGAA |
| CadO_V1_FWD | CATACTCAAATGAACATTTGAGTGAG |
| CadO_V2_FWD | CACACTCAAATGAACATTTGAGTAAG |
| CadO_V3_FWD | CATACTCAAGTGAACATTTGAGTAAG |
| CadO_V4_FWD | CACACTCAAATGAACATTTGAGTGAG |
| CadO_V5_FWD | CACACTCAAATGAACGTTTGAGTGAG |
| CadO_V6_FWD | CACACTCAAACGAACATTTGAGTGAG |
| CadO_V7_FWD | CACACTCAAACGAACGTTTGAGTGAG |
| CadO_V8_FWD | CACACTCAAGTGAACACTTGAGTGAG |
| CadO_V9_FWD | CACACTCAAACGAACGCTTGAGTGAG |
| CadO_V10_FWD | CACACTCAAGCGAACGTTTGAGTGAG |
| CadO_V11_FWD | CACACTCAAGTGAACGCTTGAGTGAG |
| CadO_V12_FWD | CACACTCAAGCGAACACTTGAGTGAG |
| CadO_V13_FWD | CACACTCAAGCGAACGCTTGAGTGAG |
| CadO_WT_REV | TTCATTCAAATATTTATTTGAGTGTA |
| CadO_V1_REV | CTCACTCAAATGTTCATTTGAGTATG |
| CadO_V2_REV | CTTACTCAAATGTTCATTTGAGTGTG |
| CadO_V3_REV | CTTACTCAAATGTTCACTTGAGTATG |
| CadO_V4_REV | CTCACTCAAATGTTCATTTGAGTGTG |
| CadO_V5_REV | CTCACTCAAACGTTCATTTGAGTGTG |
| CadO_V6_REV | CTCACTCAAATGTTCGTTTGAGTGTG |
| CadO_V7_REV | CTCACTCAAACGTTCGTTTGAGTGTG |
| CadO_V8_REV | CTCACTCAAGTGTTCACTTGAGTGTG |
| CadO_V9_REV | CTCACTCAAGCGTTCGTTTGAGTGTG |
| CadO_V10_REV | CTCACTCAAACGTTCGCTTGAGTGTG |
| CadO_V11_REV | CTCACTCAAGCGTTCACTTGAGTGTG |
| CadO_V12_REV | CTCACTCAAGTGTTCGCTTGAGTGTG |
| CadO_V13_REV | CTCACTCAAGCGTTCGCTTGAGTGTG |
